# An Injectable Antisense Oligonucleotide Approach to Treat Obesity in Preclinical Models

**DOI:** 10.64898/2026.10.12.753973

**Authors:** Simon T Bond, Michael F Keating, Yingying Liu, Christine Yang, Shannen M Walker, Aaron W Jurrjens, Kevin H Liu, Adam C Parslow, Artika P Nath, Sergio Ruiz Carmona, Michael Inouye, Anna C Calkin, Brian G Drew

## Abstract

Obesity leads to metabolic dysregulation and thus drives complications including type 2 diabetes, cardiovascular disease, steatotic liver disease and some cancers. Obesity is primarily driven by lifestyle factors, which impacts individuals differently depending on their genetic susceptibility. Studies on this genetic risk have provided major insights into gene networks that modulate obesity development, including incretins and their regulatory circuits. Indeed, incretin-based therapeutics (i.e. GLP1 RA) have proven to be enormously successful in promoting weight loss, but do so with several ongoing challenges including significant weight rebound upon cessation, loss of lean mass, and side effects that reduce therapeutic longevity. Thus, ongoing research is necessary to identify additional interventions that overcome or minimise these concerns. To this end, in our previous work we have identified a novel lipid regulatory protein with potential to influence lipid storage. In the current study, we use antisense oligonucleotides (ASO) to silence this protein in mouse models of diet induced obesity, and demonstrate that once weekly IP injections of ASO prevents the development of obesity, and defends against the progression of existing obesity. This is shown to be effective in multiple mouse strains fed different diets over various time frames, with gene targeting ASOs leading to reductions in fat mass, no change in lean mass, along with improvements in whole body glucose tolerance and steatotic liver disease. Additional studies demonstrate that this phenotype is likely driven by effects in adipose tissue, but not liver, and is not linked to activation of the Type 1 Interferon response, a previously implicated off target effect of ASO administration. Thus, these studies demonstrate that our injectable ASO approach in mice can successfully treat obesity in the absence of major change in lean mass, thus positioning this intervention as a therapeutic program that works alternatively to, or in conjunction with current weight loss medicines.

## INTRODUCTION

Obesity is a chronic metabolic disease characterised by an excessive accumulation of adipose tissue that subsequently disrupts systemic energy homeostasis and increases the risk of numerous comorbidities^1^. The global prevalence of obesity has risen dramatically over recent decades, largely driven by changes in diet, physical activity, and broader social and environmental factors^1^. Despite these strong lifestyle and environmental influences, inherited susceptibility to obesity varies considerably between individuals, highlighting genetics as an important factor influencing the development and progression of obesity and its associated comorbidities. Multiple large-scale human and mammalian systems genetics approaches have now identified numerous genes and networks that are associated with obesity susceptibility^2^. Subsequent functional studies have also demonstrated that these obesity-associated genes participate across a diverse range of biological pathways that govern adiposity through effects on appetite and feeding behaviour, energy expenditure, lipid metabolism, nutrient utilisation, and endocrine signalling^2,3^.

Importantly, these discoveries have expanded our understanding of the biological pathways that regulate body weight and adiposity, and have established a framework for the identification of novel therapeutic targets. The recent success of glucagon-like peptide-1 (GLP-1) based therapies, and related incretin receptor agonists, has further demonstrated the clinical value of targeting key pathways involved in energy balance and metabolic homeostasis^4^. However, a number of limitations remain with current obesity therapeutics, including reduced efficacy in some individuals, adverse effects including loss of muscle mass, and the tendency for weight regain following cessation of therapy^5,6^. Consequently, there remains a need to identify novel therapeutic targets that can either enhance or compliment the efficacy of existing incretin-based therapies, or that can provide alternative treatment strategies through distinct mechanisms.

In previous work, we have used a systems genetics approach in mice from the hybrid mouse diversity panel (HMDP, >100 genetically diverse strains) to identify 26S Proteasome Non-ATPase Regulatory Subunit 9 (PSMD9) as a novel regulator of lipid metabolism and storage^7^. PSMD9 is a multifunctional protein within the 26S proteasome which has been implicated in diverse cellular processes including protein degradation, transcriptional regulation, and metabolic control^8,9^. Genetic and preclinical studies have demonstrated that altered PSMD9 expression influences lipid storage, adiposity and metabolic homeostasis, suggesting that PSMD9 may represent a promising target for obesity^7^. Moreover, data from large human genetic resources (Common Metabolic Diseases Knowledge Portal) demonstrate that SNPs in the PSMD9 locus are strongly associated with body weight and BMI. However, to our knowledge there has been no direct experimental evidence to demonstrate whether manipulating PSMD9 abundance can impact on obesity related phenotypes.

In the present study, we investigated whether silencing of PSMD9 using anti-sense oligonucleotides (ASOs) could prevent the development of obesity, or treat previously established obesity. Using multiple mouse strains (C57BL/6J and DBA/2J), diet interventions, and tissue targeting strategies, we evaluated the effects of PSMD9-ASOs on body weight (including fat mass and lean mass), glucose homeostasis, and liver pathology. Our findings identify PSMD9-ASOs as a promising approach for treating obesity and its complications including robust protection against diet-induced weight gain, improved metabolic health, and enhanced weight loss responses during dietary intervention. Together, these findings establish this approach as a therapeutic modality that may work alternatively to, or in conjunction with current weight loss medicines.

## RESULTS

### Treatment with ASOs Targeting PSMD9 Protects Against the Development of Obesity

In previous work from our group, we have designed, optimised and validated several antisense oligonucleotides (ASOs) that target PSMD9 with no identified health decrements in mice^7^. From this work we identified a native ASO (non-tissue targeted) that when delivered to mice once weekly (25mg/kg) by intraperitoneal (IP) injection for 4-weeks, led to complete ablation of PSMD9 in several tissues^7^, and demonstrated reduced liver steatosis in mouse models fed a high fat and high cholesterol diet (Western Diet)^7^. In the present work, we set out to undertake longer term high fat feeding studies, which are more appropriate for investigating the chronic effects on development of metabolic pathologies including Metabolic dysfunction Associated Steatohepatitis (MASH - formerly NASH).

For these experiments, we employed the Amylin Liver NASH (AMLN) diet model, which induces obesity, insulin resistance, and hepatic steatosis, inflammation, and fibrosis, representing many of the key features of human MASH^10,11^. Using this model, we investigated the impact of PSMD9-ASOs on these phenotypes. Specifically, mice were fed an AMLN diet (high in fat, cholesterol and fructose) for 24-weeks, and were administered weekly IP injections over 24-weeks with either native PSMD9-ASO, native Scrambled ASO (Scr-ASO), or Vehicle (Saline) **(Supplementary Figure S1A)**. We demonstrate in this model that once weekly IP injections with native PSMD9-ASO (25mg/kg) substantially prevented weight gain over 24-weeks of diet-intervention **(Figure 1A),** compared with animals that received the same diet but were treated with either saline or scrambled ASO (Scr-ASO). This prevention in weight gain became significant from approximately 7-8 weeks after the first injection **(Figure 1A)**, confirming our previous observation of no change in body weight after 4-weeks of ASO treatment in C57BL/6J mice^7^. This prevention in weight gain was driven primarily by a lack of accretion in fat mass (**Figure 1B & Supplementary Figures S1B)**, with no obvious negative impact on lean mass (**Figure 1C & Supplementary Figures S1C)** and thus a dramatic improvement in the fat to lean mass ratio **(Supplementary Figure S1D)**. This was shown to impact both visceral and subcutaneous fat pads to a similar extent, with significant reductions in both gonadal (G.Adipose: visceral) and inguinal (I.Adipose: subcutaneous) adipose depot weights at the end of the study **(Figure 1D)**. There was no significant change in the weight of other organs including heart, brown adipose (B.Adipose) and skeletal muscle (Quadricep), although the livers of PSMD9-ASO treated mice were significantly heavier (**Figure 1D)**. These PSMD9-ASO induced improvements in adiposity resulted in an improved glucose tolerance at the end of the study as measured by oral glucose tolerance test (oGTT) (**Figures 1E & 1F)**.

**Figure 1:**
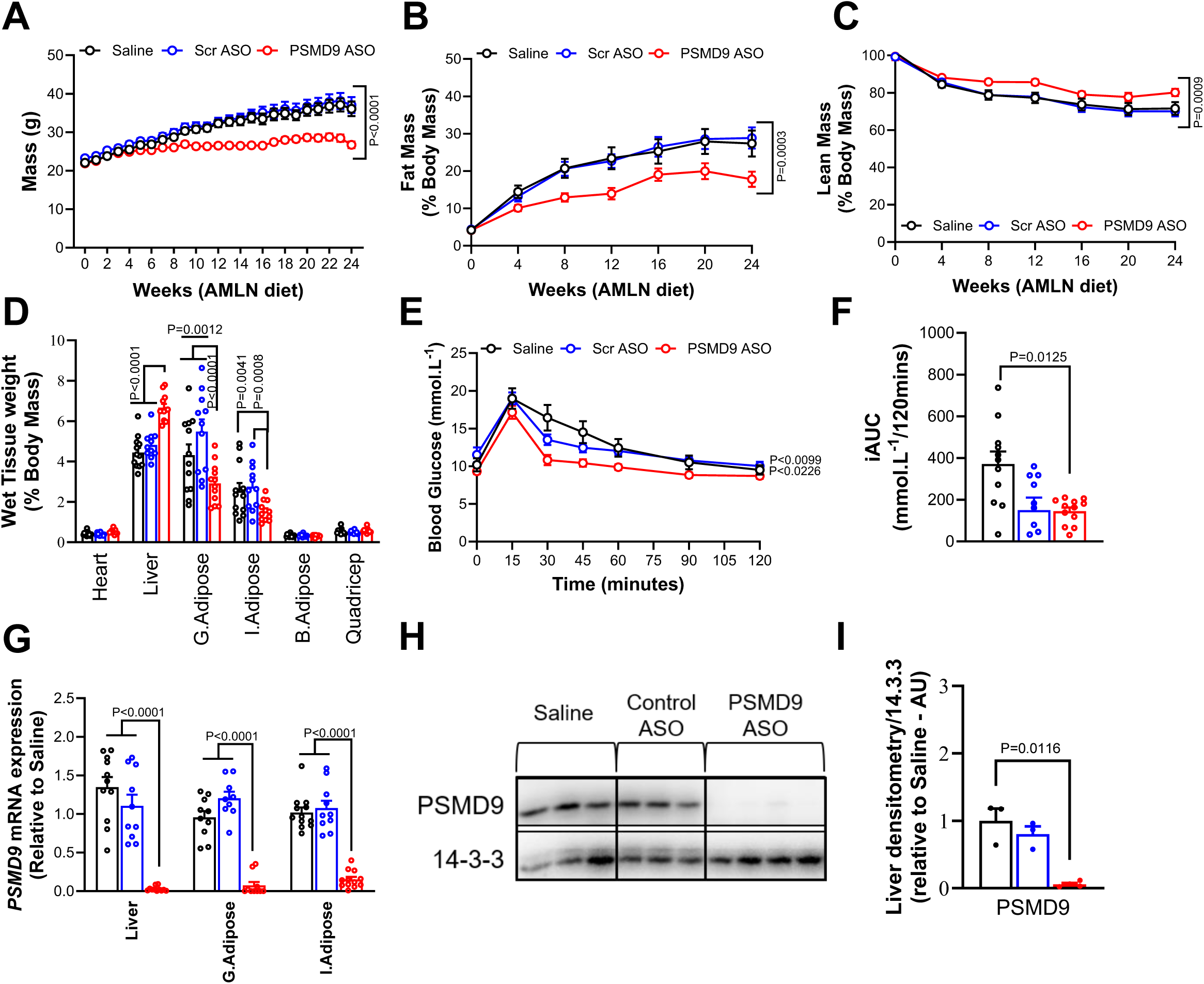
Treatment with ASOs Targeting PSMD9 Protects Against the Development of Obesity. **(A)** Weekly whole-body weights in C57BL/6J mice treated with either Saline (black, n=12), Scrambled ASO (blue, n=11), or PSMD9-ASO (red, n=12). **(B)** Progressive percent fat mass, **(C)** percent lean mass and **(D)** percent wet tissue mass of Saline, Scrambled ASO and PSMD9-ASO treated C57BL/6J mice as measured by percent total body mass. **(E)** Glucose tolerance test performed in Saline, Scrambled ASO and PSMD9-ASO treated C57BL/6J mice measuring change in plasma glucose as measured every 15 minutes over the 2 hours post-glucose gavage, with **(F)** corresponding incremental area under the curve analysis. **(G)** *Psmd9* mRNA expression in Saline, Scrambled ASO and PSMD9-ASO treated C57BL/6J mice liver, visceral adipose tissue (gonadal) and subcutaneous adipose tissue (inguinal). **(H)** Immunoblot for PSMD9 in liver from Saline, Scrambled ASO and PSMD9-ASO treated C57BL/6J mice with **(I)** corresponding PSMD9 protein quantification normalised to 14-3-3. Data are presented as mean ± SEM. P-value determined with correction for multiple comparisons by two-way ANOVA (panels A-C, D-E, & G), or one-way ANOVA (panels F & I).

Although we have demonstrated in our prior work that PSMD9-ASOs effectively reduced PSMD9 mRNA and protein abundance in several tissues in short-term studies^7^, we wanted to confirm similar effects on PSMD9 silencing in mice treated for 24-weeks. Accordingly, we performed qPCR to investigate the abundance of PSMD9 mRNA in liver, gonadal adipose tissue (G.Adipose) and inguinal adipose tissue (I.Adipose). These data demonstrate almost complete ablation of PSMD9 in C57BL/6J mice **(Figure 1G)**. Importantly, there was no change in PSMD9 abundance with either saline or Scr-ASO treatment. Western blotting for PSMD9 abundance in liver also demonstrated an almost complete loss of PSMD9 protein in PSMD9-ASO treated mice (**Figures 1H & 1I)**, confirming that the ASOs were successful in targeting and silencing PSMD9.

In order to demonstrate that the effect on adiposity and obesity complications was not a phenomenon specific to the C57BL/6J strain, we also repeated the vast majority of these experiments in a second strain of mice, DBA/2J. Consistently, these data show that once weekly IP injection of PSMD9-ASOs in DBA-2J mice led to a similar reduction in body weight, adiposity and fat:lean mass ratio as shown through various readouts **(Supplementary Fig S1E – S1K)**. These data provide strong evidence that once weekly injections of ASOs targeting PSMD9, lead to long term silencing of PSMD9 in multiple tissues of two independent mouse strains, which is sufficient to prevent weight gain and reduce adiposity in a long-term model of diet induced metabolic dysregulation. Subsequent work was thus undertaken to focus on the impact of PSMD9-ASOs on obesity related complications, and whether the treatment is effective at reversing or slowing the progression of established obesity.

### Treatment with whole body PSMD9-ASOs Reduced the Severity of Hepatic Steatosis

Given the demonstrated impacts on body weight and adiposity, we were interested in firstly, understanding the potential mechanism for this weight loss, but also the impacts this weight loss has on tissue complications including hepatosteatosis. With regards to mechanisms driving weight loss, interestingly we did not observe any striking differences in the main physiological pathways that would normally be associated with changes in energy storage. This included no significant difference in food intake between groups (**Figure 2A),** nor any changes in energy expenditure (EE), respiratory quotient (RQ), EE after adjustment for body mass using ANCOVA, or activity in these mice as measured by Promethion metabolic cages **(Figures 2B-2E & Supplementary Figures S2A & S2B)**. Whilst these changes were not significantly different, there were trends to suggest that weight adjusted energy expenditure was maintained in PSMD9-ASO treated mice even though they were lighter (left shift in the plot – red dots, **Figure 2D**), and a trend for an increased activity in PSMD9-ASO (**Figure 2E**). These slight changes are too subtle to detect significant differences in 2-3 days of testing, but over several weeks would likely be sufficient to drive changes in body composition as observed. This notwithstanding, in order to show that this effect wasn’t due to toxicity of the treatment, we performed blood pathology measures at the end of the study. These data demonstrate that there were no detectable differences in various plasma readouts including total protein, albumin, globulin, creatinine or bilirubin **(Supplementary Figure S2C**). Moreover, there were no differences in plasma β-hydroxybutyrate and uric acid **(Supplementary Figure S2D**). Finally, although modestly elevated, there were no pathological changes in the liver enzyme alanine aminotransferase (ALT) (below 200IU/L) **(Supplementary Figure S2E**), with a separate measure of liver function (albumin to globulin ratio) showing no difference between groups **(Supplementary Figure S2F**). Thus overall, we attribute the difference in weight gain and adiposity in these animals to a subtle change in energy expenditure over an extended period of time, that was not due to any major effects on systemic toxicity.

**Figure 2:**
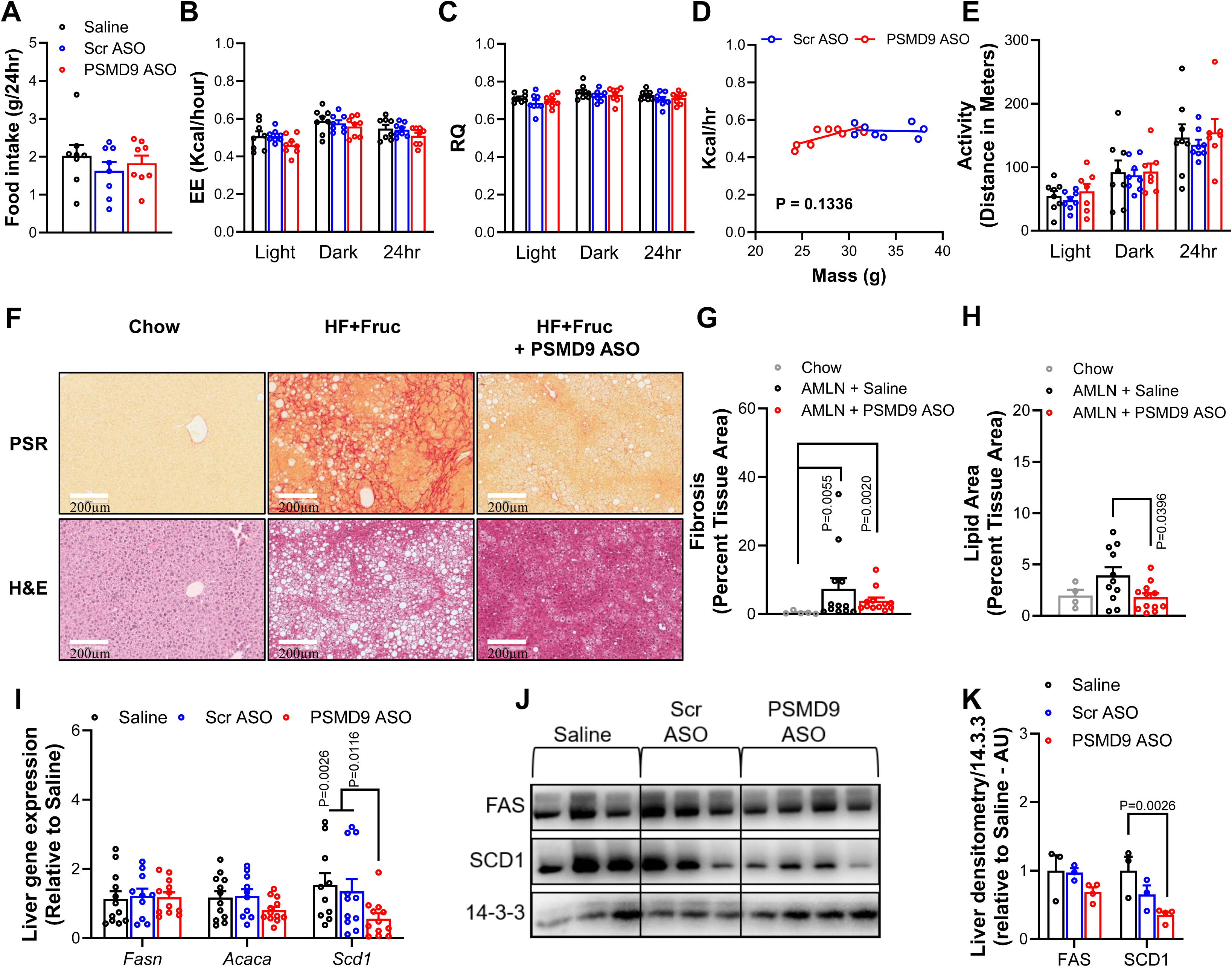
Treatment with whole body PSMD9-ASOs Reduced the Severity of Hepatic Steatosis. C57BL/6J mice treated with either Saline (black, n=12), Scrambled ASO (blue, n=11), or PSMD9-ASO (red, n=12) were placed in metabolic cages (12hrs light & 12hrs dark) to determine 24hr averages of **(A)** food intake, **(B)** energy expenditure (EE), **(C)** respiratory quotient (RQ), **(D)** EE as a measure of total body mass (ANCOVA) and **(E)** total activity. **(F)** Sections of liver from Saline, Scrambled ASO, and PSMD9-ASO treated C57BL/6J mice were analysed using PSR and H&E staining to determine **(G)** fibrosis area and **(H)** lipid area as a percent of total tissue area. **(I)** *Fasn, Acaca,* & *Scd1* gene expression and **(J)** protein expression with **(K)** corresponding protein quantification normalised to 14-3-3 measured in liver from Saline, Scrambled ASO, and PSMD9-ASO treated C57BL/6J mice. Data are presented as mean ± SEM. P-value determined with correction for multiple comparisons by two-way ANOVA (panels A-C, E, I & K), or one-way ANOVA (panels G-H), or ANCOVA (panel D).

This being the case, we next investigated the effect of PSMD9-ASO treatment on liver pathology on these mice fed an AMLN diet. As discussed above, long term AMLN diet feeding drives MASH, which is characterised by steatosis, inflammation and fibrosis in the liver, and is a major driver of cirrhosis and liver cancer in humans and rodents^12,13^. One of the most reliable and effective ways to measure MASH is by histological characterisation of liver sections. We therefore performed staining of liver sections with picrosirius red (PSR) for fibrosis, and hematoxylin and eosin (H&E) staining for analysis of morphology including steatosis (lipid droplet area). As can be observed from representative images from these analyses **(Figure 2F)**, we demonstrate that AMLN diet leads to substantial increases in the level of fibrosis in the liver compared with chow fed mice, as characterised by the appearance of red fibrils in PSR staining. The intensity of this effect is significantly decreased in mice treated with PSMD9-ASOs, as confirmed by the quantitation of the PSR intensity **(Figure 2G)**. With regards to steatosis, we observed a considerable increase in the appearance of lipid droplets in the H&E stain from chow to AMLN diet **(Figure 2H)**, which is also significantly reduced in mice fed AMLN diet but receiving PSMD9-ASO (**Figure 2H)**. Using qPCR, we demonstrate that genes relevant to lipid metabolism in the liver, including those that regulate de novo lipogenesis (DNL; *Fasn*, *Acaca*, *Scd1*), are altered in mice that were treated with PSMD9-ASO (**Figure 2I)**. Consistent with transcriptional changes in the DNL pathway, we also observe similar changes at the protein level in these pathways (particularly SCD1), as shown by western blot and quantitation of protein abundance (**Figures 2J & 2K)**

Given the beneficial effects of PSMD9-ASOs on the prevention of obesity and subsequent improvements in metabolic health and MASH, we next determined whether these same effects could be recapitulated in mice that had lived with obesity for several weeks, before receiving the PSMD9-ASO therapy. This approach has greater translational relevance, as it more closely reflects the therapeutic setting in which interventions are administered in humans, who would almost certainly already be overweight or obese before receiving an anti-obesity therapy.

### PSMD9-ASOs Improve Adiposity and Glucose Tolerance in Established Obesity

To investigate the effects of PSMD9-ASOs in established obesity, we fed mice either a HFD for 14-weeks (HFD→HFD), or a HFD for 3-weeks followed by a switch back to chow diet (HFD→Chow) for a further 11-weeks to mimic a healthy diet intervention. After 6-weeks of any diet, the mice were then commenced on weekly ASO/Vehicle treatments (Saline, Scr-ASO and PSMD9-ASO) **(Supplementary Figure S3A)**. At week 3, when animals were switched back to chow, it can be seen that body weight plateaued at ∼15% above starting weight, however those that remained on HFD increased to ∼50% above starting weight by week 6, and finished at ∼75% heavier than their starting weight by end of the study. **(Figure 3A, Supplementary Figure S3B**). Animals that received control treatments (Saline and Scr-ASO) and HFD, continued to put on weight from 6 weeks onwards, whilst mice on PSMD9-ASO on HFD were protected, ending up at ∼50% above starting weight, being half of that which was accumulated by mice receiving the control treatments. For the animals that were treated with both a diet change (switch to chow) and PSMD9-ASO, the ASOs had an additive effect on reducing body weight, reducing the final end weight almost back to their starting weight, where Scr-ASO treated mice finished ∼20% above starting weight **(Figure 3A & Supplementary Figure S3B**). These changes in body weight were mostly related to changes in body fat **(Figure 3B and Supplementary Figure S3C),** with no discernible reduction in lean mass **(Supplementary Figures S3D & S3E)**. These findings were further confirmed with significant reductions observed in adipose tissue depot weights, including reduced gonadal and inguinal adipose in HFD→HFD fed mice and reduced gonadal adipose in HFD→Chow fed mice **(Supplementary Figures S3F & S3G)**.

**Figure 3:**
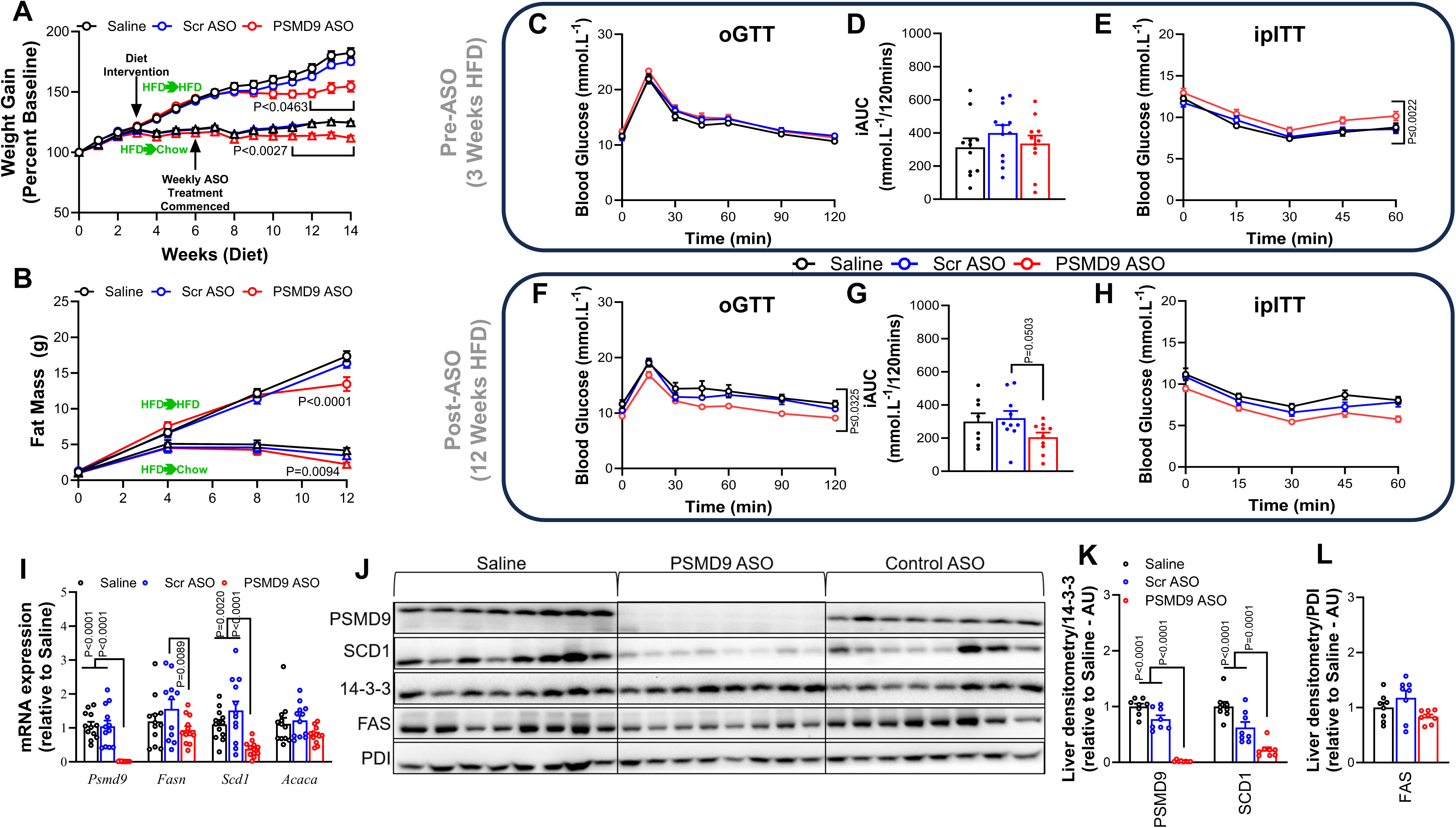
PSMD9-ASOs Improve Adiposity and Glucose Tolerance in Established Obesity. **(A)** Weekly weight gain (percent from baseline) and **(B)** progressive fat mass in C57BL/6J mice with established obesity treated with either high fat diet (HFD) + Saline (black circle, n=12), Scrambled ASO (blue circle, n=12), or PSMD9-ASO (red circle, n=12) or CHOW + Saline (black triangle, n=12), Scrambled ASO (blue triangle, n=12), or PSMD9-ASO (red triangle, n=12). Glucose (including corresponding iAUC) and insulin tolerance tests were performed in HFD + Saline, Scrambled ASO, and PSMD9-ASO treated C57BL/6J mice after **(C-E)** 3-weeks of HFD and before weekly ASO injections commenced and **(F-H)** after 12-weeks of HFD and 6-weeks ASO treatment. **(I)** *Psmd9, Fasn, Scd1* & *Acaca* gene expression and **(J)** protein expression, with corresponding **(K)** quantification to 14-3-3 and **(L)** PDI, in livers from HFD + Saline, Scrambled ASO, and PSMD9-ASO treated C57BL/6J mice. Data are presented as mean ± SEM. P-value determined with correction for multiple comparisons by two-way ANOVA (panels A-C, E-F, & H-I), or one-way ANOVA (panels D, G, K & L).

Similar to our studies earlier in which we prevented the development of obesity with PSMD9-ASOs, in these experiments of established obesity, we again observed no significant differences in energy expenditure (EE), respiratory quotient (RQ), EE after adjustment for body mass using ANCOVA, food intake, or activity in these mice as measured by Promethion metabolic cages **(Supplementary Figures 3H-3L)**. These data further suggest that even though there were noticeable differences in body weight, including reductions in fat mass, that any changes in metabolic activity were too subtle to detect in short-term analysis using the Promethion metabolic cages (48-hour period).

Importantly, we did observe non-significant trend for improvements in fasting blood glucose in HFD→HFD fed mice administered PSMD9-ASO, suggesting an improved glucose homeostasis in these animals **(Supplementary Figure S3M)**. This was associated with improvements in glucose tolerance where at 3-weeks of HFD and before ASO treatment, all groups had similar tolerance to glucose and insulin **(Figures 3C-3E)**. However, after a further 9-weeks of HFD and 6-weeks of ASO treatment, the PSMD9-ASO group had significantly improved glucose tolerance compared with the Saline and Scr-ASO groups **(Figures 3F & 3G)**. Moreover, even though the PSMD9-ASO treated group had a poorer insulin tolerance at baseline, after 6-weeks of PSMD9-ASO treatment this insulin tolerance was improved compared with the Saline and Scr-ASO groups **(Figure 3H)**, consistent with the lower fasting blood glucose in these animals **(Supplementary Figure S3M)** Thus, in a model of existing obesity, PSMD9-ASO treatment led to reductions in adiposity and weight gain, while preventing declines in diet induced glucose tolerance and insulin resistance.

Similar to our earlier studies, *Psmd9* mRNA abundance was almost completely ablated in the liver in these studies, which coincided with reductions in mRNA expression of *Fasn* and *Scd1* **(Figure 3I),** and consistent with significant protein reductions in PSMD9 and SCD1, when compared with Saline and Scr-ASO groups **(Figures 3J-3L)**.

Given these consistent findings in the liver tissue of mice treated with PSMD9-ASO in both prevention and regression studies thus far, including changes in DNL and improvements in whole body glucose metabolism, we suspected that the liver was likely the important tissue driving these metabolic changes. Accordingly, to test this directly, we next generated liver targeted PSMD9-ASOs (GalNAc conjugated ASOs), and repeated the “prevention” studies using the 24-week AMLN design in C57BL/6J mice.

### Liver Targeted PSMD9-ASOs do not Impact Body Weight, Adiposity or Metabolic Health

To undertake liver targeting PSMD9-ASOs experiments, we used GalNAc conjugated ASOs. This approach has been demonstrated in several studies previously to allow for ASGR dependent uptake of ASOs in hepatocytes. For these studies we used the same ASO and sequence for both scrambled (Scr) and PSMD9-ASOs to the previous experiments, with the exception being that they were GalNAc conjugated. For these experiments we fed C57BL/6J mice an AMLN diet for 24-weeks in conjunction with weekly GalNAc ASO injections (1mg/kg; Saline, Scr GalNAc, PSMD9 GalNAc) **(Figure 4A)**. Firstly, to confirm GalNAc PSMD9-ASO tissue specificity, we immunoblotted liver and adipose tissue, demonstrating PSMD9 silencing in liver, but not in adipose tissue **(Figure 4B)**. However, to our surprise, after 24-weeks of AMLN diet and GalNAc ASO treatment, we observed no noticeable reductions in whole-body weight between groups **(Figure 4C)**. In addition, GalNAc PSMD9-ASO treatment did not reduce fat mass or lean mass **(Figures 4D-4G)** compared to both saline and GalNAc Scr-ASO. This was also confirmed in tissue weights at the end of the study, with no reductions in adipose weight, but a slight increase in liver weight of GalNAc PSMD9-ASO treated animals **(Figure 4H)**. There were no significant changes in food intake or glucose tolerance between all groups **(Supplementary Figures S4A-S4C)**. Moreover, this was not a strain specific effect, because there were almost identical data generated when we repeated the GalNAc PSMD9-ASO experiments in the DBA/2J strain **(Supplementary Figures 4D & 4E)**. Given this lack of effect with liver specific targeting of PSMD9-ASOs, we next turned our attention to adipose tissue, which could also be responsible for major changes in metabolic health and adiposity.

**Figure 4:**
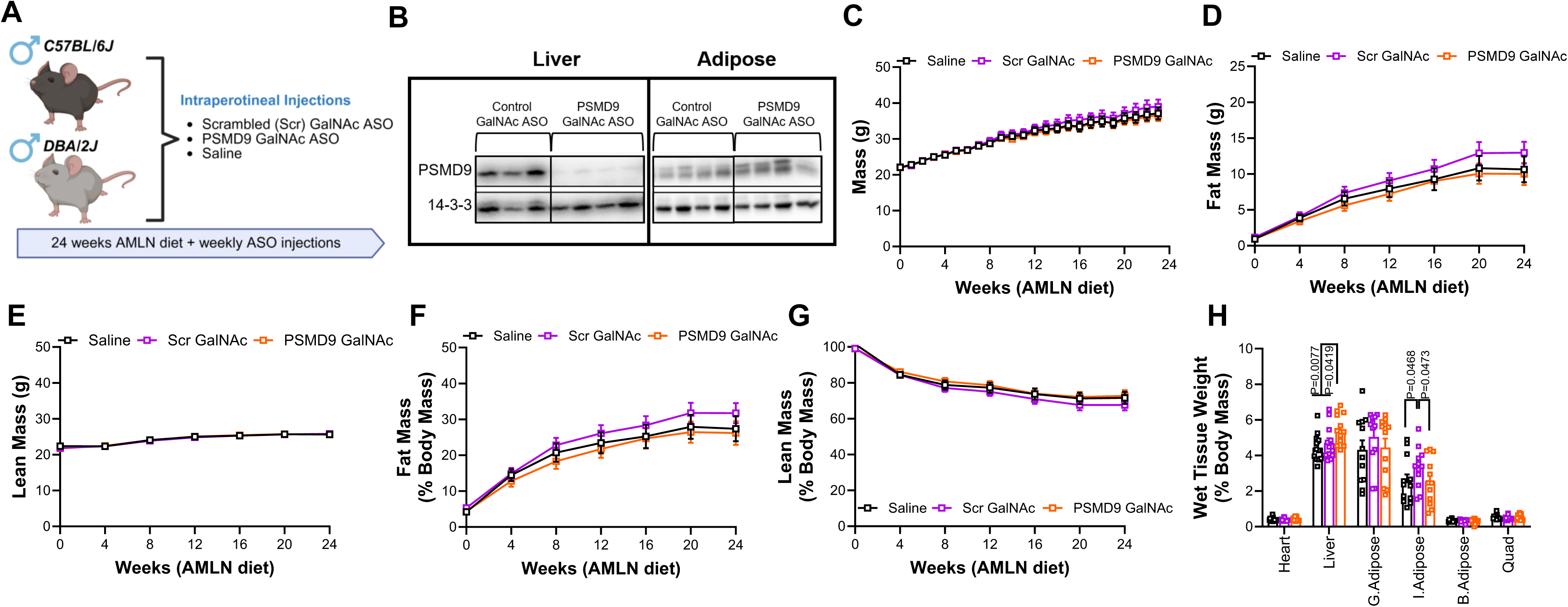
Liver Targeted PSMD9-ASOs do not Impact Body Weight, Adiposity or Metabolic Health. **(A)** Schematic outlining animal study to investigate effect of weekly liver targeted PSMD9 GalNAc ASO administration in C57BL/6J and DBA/2J mice. **(B)** Immunoblot for PSMD9 in liver from Saline, Scrambled GalNAc ASO and PSMD9 GalNAc ASO treated C57BL/6J mice. **(C)** Weekly body weight, **(D)** monthly fat mass, **(E)** monthly lean mass, **(F)** monthly percent fat mass, and **(G)** monthly percent lean mass in C57BL/6J mice treated with either Saline (black, n=12), Scrambled GalNAc ASO (purple, n=12) and PSMD9 GalNAc ASO (orange, n=12). **(H)** Wet tissue weights as a percent of total body mass from C57BL/6J mice treated with GalNAc ASOs. Data are presented as mean ± SEM. P-value determined with correction for multiple comparisons by two-way ANOVA (panels C-H).

### Native PSMD9-ASOs Impart Major Impacts on Adipose Tissue

Having identified that liver specific PSMD9-ASOs did not recapitulate the adiposity effects observed with ASOs that target all tissues (native ASOs), we went back to our original studies and investigated the effects of native PSMD9-ASOs in adipose tissue. Similar to gene expression in the liver, we confirmed by qPCR analysis that *Psmd9* mRNA abundance was reduced in adipose, with a similar reduction observed in DNL related genes including *Fasn* and *Scd1* that was observed in the livers, but no difference in *Acaca* expression **(Supplementary Figure S5A)**. Immunoblotting further confirmed that PSMD9 was depleted in adipose tissue, but interestingly, there were no significant changes in the lipolytic state, as assessed by HSL phosphorylation at Ser563 **(Figures 5A & 5B)** suggesting that lipolysis was an unlikely pathway involved in the phenotype.

**Figure 5:**
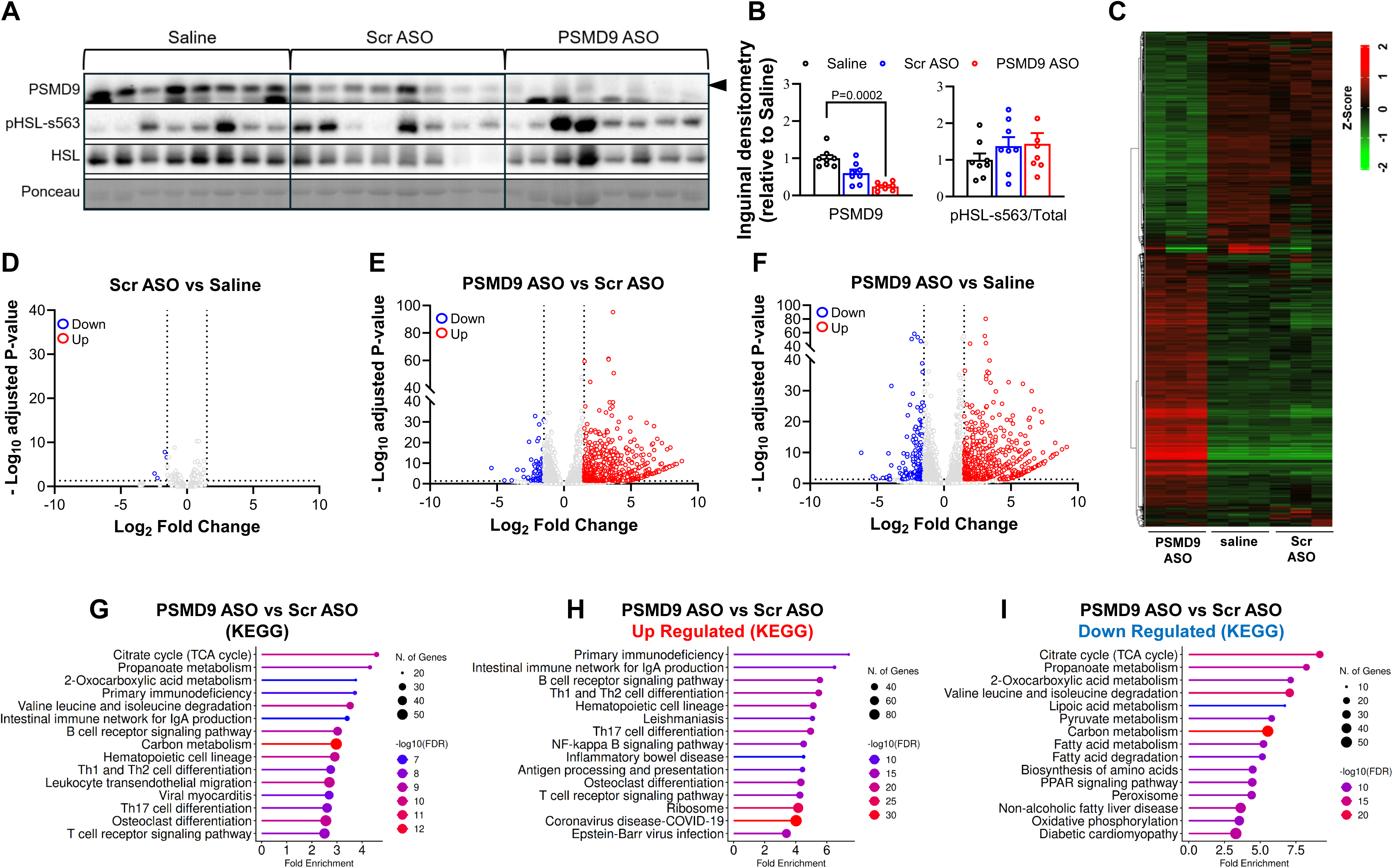
Native PSMD9-ASOs Impart Major Impacts on Adipose Tissue. **(A)** Immunoblot for PSMD9 in gonadal adipose tissue from C57BL/6J mice with established obesity and treated with either Saline, Scrambled ASO or PSMD9-ASO after 6-weeks of HFD for 8-weeks (regression cohort from figure 3), with **(B)** corresponding protein quantification. From the same mice, transcriptomic analysis of white adipose tissue was performed to measure RNA abundance. Computational analysis of the data was performed including **(C)** hierarchical clustering and volcano plots of differentially expressed genes (DEGs) comparing **(D)** Scr-ASO with Saline, **(E)** PSMD9-ASO with Scr-ASO **(F)** and PSMD9-ASO with Saline (blue=down regulated genes, red=upregulated genes). Pathway enrichment analysis (KEGG) on DEGs from PSMD9-ASO vs Scr-ASO depicting **(G)** up and down regulated enriched pathways, **(H)** pathways that were significantly upregulated between the genotypes and **(I)** pathways that were significantly downregulated between the genotypes. Data are presented as mean ± SEM, with P-value determined with correction for multiple comparisons by one-way ANOVA (panel B). Analyses of transcriptomic data (C-I) were corrected using Benjamini-Hochberg FDR, n=3/group with DEGs determined by adj P-value < 0.05 and FDR > 1.5.

Given these robust impacts of PSMD9-ASO on adipose tissue, we next performed RNA sequencing on adipose tissue (n=3/group) to get a broader perspective of the pathways that were being modulated by our treatment. Across all three groups we first demonstrated that there was consistency in overall gene counts **(Supplementary Figure S5B)**, and that hierarchical clustering **(Figures 5C)** and PCA analysis **(Supplementary Figure S5C)** demonstrated that there is significant variance in the transcriptome between the PSMD9-ASO and both the Scr-ASO and vehicle (saline) control groups, with negligible expression differences between the two control groups (demonstrating no effect of the Scr-ASO). This is further demonstrated through differential gene expression analyses as shown in volcano plots with FDR<0.05 and FC>1.5 (red dots = increased expression, blue dots = reduced expression for each comparison, **Figures 5D-5F)**. The number of DEGs between groups was only notable in PSMD9-ASO groups compared to either control **(Supplementary Figure S5D)**, with only 4 DEGs different between the two control groups (4 down, 0 up), but 779 DEGs between PSMD9 and Scr-ASO treated mice (84 down, 695 up), and 852 DEGs between PSMD9-ASO and Saline treated mice (194 down, 658 up). Pathway analysis performed on DEGs between PSMD9-ASO and Scr-ASO treated mice demonstrated an enrichment in metabolic and immune responsive pathways **(Figure 5G)**, in particular an enrichment of immune related pathways in upregulated genes **(Figure 5H)**, and an enrichment in metabolic related pathways in downregulated genes **(Figure 5I)**.

### Native PSMD9-ASOs Protect from Obesity in IFNAR-KO Mice

This immune-centric phenotype observed in the adipose RNA-seq piqued our interest. Whilst this could be an important part of our phenotype relating to adipose tissue remodelling, it could also be related to previous work that has reported that certain ASO compositions could initiate the type 1 interferon response (T1IFN) in adipose tissue of mice, and promote reductions in fat mass^14^. Therefore, to discount the possibility that our PSMD9-ASO was inducing weight loss in a T1IFN dependent manner, we performed a further set of animal studies. Specifically, we repeated our PSMD9-ASO studies in mice lacking an intact T1IFN response using the IFNAR1 KO (IFNAR-KO) mouse^15^, and fed them a HFD to promote obesity. To do this, IFNAR-WT and IFNAR-KO littermates were generated and administered a HFD in conjunction with weekly native ASO injections (PSMD9-ASO, Scr-ASO and Saline) for 17-weeks **(Figure 6A)**. Firstly, it was noted that IFNAR-KO mice were smaller and had a lower body mass than IFNAR-WT mice (**Supplementary Figures S6A)**, consistent with previous studies demonstrating these mice are partially protected from HFD induced obesity^16^. Therefore, for us to examine the effects of our PSMD9-ASO treatment, we focussed on increased weight gain from baseline. Importantly, and consistent with our previous models, this data demonstrate that IFNAR-KO mice fed a HFD and administered PSMD9-ASOs, are protected from increased body weight compared with IFNAR-KO mice administered control treatments **(Figure 6B and Supplementary Figures S6A & S6B)**. This was shown to be related to reductions in fat mass **(Figure 6C** and **Supplementary Figure S6C)**, with modest to no effects on lean mass **(Supplementary Figures S6D & S6E),** all of which was confirmed in wet tissue weight analysis **(Supplementary Figure S6F)**. Thus, despite the absence of a global T1IFN response, PSMD9-ASO treatment still reduced fat mass in IFNAR KO mice. This was accompanied with similar beneficial effects on glucose tolerance, with IFNAR-KO mice treated with PSMD9-ASO demonstrating improved glucose tolerance compared with IFNAR-KO mice treated with saline and Scr-ASO **(Supplementary Figures S6G & S6H)**.

**Figure 6:**
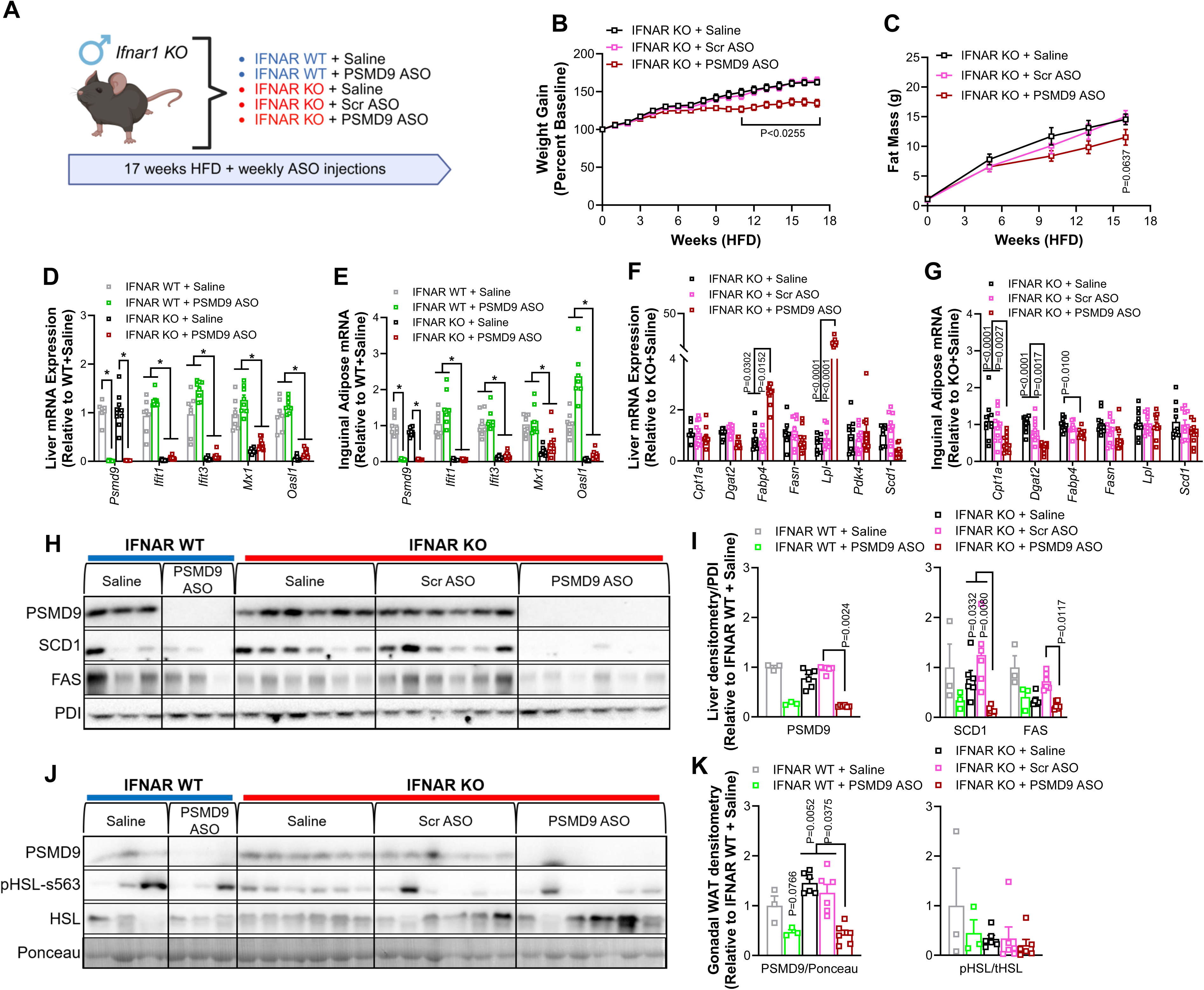
Native PSMD9-ASOs Protect from Obesity in IFNAR-KO Mice. **(A)** Schematic outlining animal study to investigate effect of weekly PSMD9-ASO administration for 17-weeks in mice fed a high fat diet (HFD) and lacking the type 1 interferon response (T1IFN - IFNAR KO). **(B)** Weekly weight gain (percent from baseline) and **(C)** progressive fat mass in IFNAR KO mice fed a HFD and treated with either Saline (black square, n=10), Scrambled ASO (pink square, n=10), or PSMD9-ASO (red square, n=11). *Psmd9* and T1IFN response gene expression in **(D)** liver and **(E)** sub-cutaneous (inguinal) adipose tissue from IFNAR WT mice fed a HFD and treated with either Saline (grey square, n=7) or PSMD9-ASO (green square, n=8) and IFNAR KO mice fed a HFD and treated with either Saline (black square, n=10) or PSMD9-ASO (red square, n=10). Metabolic gene expression in **(F)** liver and **(G)** sub-cutaneous (inguinal) adipose tissue from IFNAR KO mice fed a HFD and treated with either Saline (black square, n=9), Scrambled ASO (pink square, n=10), or PSMD9-ASO (red square, n=11). Immunoblot in IFNAR WT and KO mice used for gene expression analysis above (D-G) in **(H)** liver, with **(I)** corresponding protein quantification, and in **(J)** inguinal adipose tissue with **(K)** corresponding protein quantification. Data are presented as mean ± SEM. P-value (* denotes P<0.05) determined with correction for multiple comparisons by two-way ANOVA (panels B-G) or one-way ANOVA (panels I & K).

Given we observed similar results in the IFNAR KO mice to our previous models, we were confident that the T1INF response was not responsible for eliciting weight loss effects associated with our PSMD9-ASO. This being the case, it was also important to demonstrate that our PSMD9-ASO was still working similarly in this model, and that indeed the T1INF response was ablated in these mice.

To confirm that the PSMD9-ASO worked similarly in IFNAR KO mice, we first confirmed using qPCR that *Psmd9* mRNA expression was reduced, which was almost completely ablated in both liver **(Figure 6D)** and adipose **(Figure 6E)** of IFNAR**-**WT and IFNAR-KO mice. Moreover, these data also confirm that critical components of the T1IFN signalling response were also ablated in IFNAR KO mice, including the expression of key T1INF response genes *Ifit*, *Ifit3*, *Mx1* and *Oasl1* **(Figure 6D & 6E)**. Additional gene expression in liver **(Figure 6F)** and inguinal adipose **(Figure 6G)** further demonstrated metabolic pathways modulated by PSMD9-ASO in the absence of T1IFN response, including genes important for DNL and lipid metabolism. These data are confirmed through immunoblotting with reductions in PSMD9 in liver **(Figure 6H & 6I)** and in adipose **(Figure 6J & 6K)**. Consistent with data in C57BL/6J mice, reductions in SCD1 and FAS protein levels in the liver were also observed following PSMD9-ASO treatment in IFNAR-KO mice. Collectively, these data provide compelling evidence that PSMD9-ASO treatment had effectively identical responses in animals lacking a functional T1INF response (IFNAR-KO) in both liver and adipose tissues, to that of IFNAR-WT and C57BL/6J.mice This included consistent effects in reducing body weight and adiposity, and in altering aspects of the DNL pathway in the liver and adipose tissue. Accordingly, we conclude that the phenotypic effects of our PSMD9-ASO are not related to activation of the T1INF response.

## Conclusions

Overall, our results present compelling evidence in 8 independent mouse models across multiple strains, genetic mouse models and diet interventions, that PSMD9-ASO treatment is able to protect from the development of obesity, and to protect from complications associated with established obesity. This occurs in the absence of major changes in lean mass. The mechanism of action is unlikely to be a direct effect on the liver, but likely a peripheral effect involving the adipose tissue. Specifically, the PSMD9-ASO treatment induces profound changes in lipid metabolism and subsequent tissue remodelling in the adipose tissue compartment that is unrelated to the T1INF response, with this remodelling of adipose tissue lipid metabolism likely explaining in part the weight loss effect. Such findings highlight this therapeutic approach as a promising new avenue for continued investigation as a treatment for obesity, that may act independently or in concert with existing therapies such as incretin-based medicines.

## DISCUSSION

Obesity remains one of the leading drivers of metabolic disease worldwide. Major breakthroughs have been achieved in the past 5-10 years in the treatment of obesity, particularly with the advent of incretin-based drugs such as GLP1/GIP1-RAs. Whilst these drugs are enormously successful and have demonstrated profound impacts on reducing adiposity in millions of people, there remains increasing concerns relating to the concomitant effect of these drugs on clinically relevant reductions in lean mass, as well as a major portion of individuals not being able to tolerate these drugs for more than 1 year. Cessation of these drugs results in an almost complete rebound in fat mass after 2 years^17,18^. The combination of reduced lean mass, together with the rebound in fat mass is likely to ameliorate much of the health benefit achieved whilst taking the drugs, and has been shown for CVD^5^. Nonetheless, these major advances in obesity therapeutics should not be discounted due to these concerns, but instead should inspire the research community to continue to innovate in this space, and design optimised and alternative obesity therapeutics that improve patient outcomes.

Accordingly, in the present study we provide compelling evidence in mice for an alternative strategy to treat obesity, which involves injectable ASO therapeutic targeting of a novel lipid regulatory protein PSMD9. We demonstrate that PSMD9-ASOs provide robust protection against diet-induced obesity across multiple mouse models while simultaneously improving glucose tolerance and reducing the severity of metabolic dysfunction-associated steatohepatitis (MASH). Importantly, these PSMD9-ASOs were effective at not only preventing obesity development, but also in limiting the progression of established obesity, including further weight loss in conjunction with dietary intervention and improved glucose tolerance. Furthermore, by combining tissue-specific targeting approaches using GalNAc conjugation, and genetic deletion of the type I interferon receptor, we demonstrate that these metabolic benefits are independent of the liver, and do not involve activation of the T1IFN. Collectively, these findings establish this ASO mediated approach as a promising strategy for the development of novel medicines to treat obesity in humans.

Our previous work, using a systems genetics approach, identified PSMD9 as a candidate regulator of lipid metabolism, where we demonstrated that short-term PSMD9-ASO treatment reduced hepatic lipid accumulation^7^. However, whether this approach represented a viable therapeutic target for obesity remained unknown. By extending these studies in the current work into longer-term dietary mouse models of obesity and MASH, we now demonstrate that prolonged treatment with PSMD9-ASOs profoundly limits adipose tissue expansion. The consistency of these findings across two genetically distinct mouse strains (C57BL/6J & DBA/2J), multiple dietary paradigms (Chow, HFD, AMLN), and including both prevention and intervention studies; strongly suggests that treatment with PSMD9-ASOs regulates fundamental pathways controlling adiposity. Further to this point, one of the most striking observations was that reductions in body weight were almost entirely attributable to decreased fat mass, whilst lean mass was largely preserved. Preservation of lean tissue is an increasingly recognised limitation of current incretin-based therapies, including GLP-1 receptor agonists, where reductions in skeletal muscle contribute substantially to total weight loss^19,20^. Maintenance of lean mass is critical for preserving metabolic health, physical function and long-term weight maintenance^21^. Consequently, the selective reduction in adiposity observed following PSMD9-ASO treatment suggests that this pathway may provide a mechanistically distinct therapeutic strategy that could either be used independently or in conjunction with existing pharmacological approaches.

Interestingly, despite marked reductions in adiposity, we observed no detectable differences in food intake, whole-body energy expenditure, or activity, whilst the mice were housed in metabolic cages. Although these findings initially appear paradoxical, they are consistent with the possibility that relatively small but sustained alterations in nutrient partitioning or lipid turnover could be accumulating over prolonged treatment periods that eventually culminate in marked differences in fat mass. Alternatively, PSMD9-ASOs may influence adipocyte lipid storage efficiency, triglyceride cycling, lipolysis, or futile substrate utilisation without substantially altering whole-body oxygen consumption. Future isotope tracer studies measuring lipid flux, de novo lipogenesis and fatty acid oxidation will be important for defining the precise metabolic processes regulated by this treatment.

A significant metabolic benefit of PSMD9-ASOs was the improved glucose handling in both preventative and intervention studies. These improvements in glucose tolerance following PSMD9-ASO treatment are likely secondary to reduced adiposity and subsequent improvements in liver and muscle insulin sensitivity, although this was not investigated directly. However, expansion of adipose tissue promotes insulin resistance and accordingly, limiting adipose expansion would be expected to improve insulin sensitivity. Whether PSMD9-ASOs directly impact insulin signalling in metabolic tissues is outside the scope of the present study, but remains an important question for future investigation.

An unexpected finding from this study was that liver specific PSMD9-ASO (GalNAc ASO) was insufficient to reproduce the whole-body (native PSDMD9 ASO) metabolic phenotype. Although liver targeting GalNAc ASOs effectively reduced hepatic PSMD9 expression and altered lipogenic gene expression similar to native PSMD9-ASOs, they failed to reduce body weight or adiposity. These findings suggest that the improvements in hepatic steatosis observed following whole-body PSMD9 inhibition are unlikely to arise from direct hepatic actions. Rather, reductions in liver pathology likely reflect secondary improvements resulting from decreased adiposity and reduced spill over of fats to the liver.

Together, these findings suggest that reductions in adipose tissue as the principal site mediating the positive effects of PSMD9 inhibition, but the exact mechanisms governing this reduction in adiposity remain unknown. To investigate this further we performed transcriptomic profiling in adipose tissue following PSMD9 silencing with whole-body native PSMD9-ASOs. Subsequent pathway analysis identified that many metabolic pathways were suppressed and that immune-related pathways were potentially increased. Adipose tissue undergoes extensive immune remodelling during obesity, and interactions between adipocytes and resident immune cells play critical roles in regulating tissue expansion and remodelling^22,23^. Whether these transcriptional changes reflect altered immune cell composition or intrinsic changes within adipocytes remains to be determined.

Due to the alterations observed in immune pathways, a potential concern was ASO initiated activation of innate immune pathways. Specifically, previous studies have demonstrated that certain ASO chemistries activate the type I interferon (T1INF) response, which independently impacts adiposity irrespective of the intended gene target^14^. Accordingly, we performed studies in mice deficient in T1INF signalling (lacking *Ifnar1* gene), where we demonstrate that PSMD9-ASO mediated protection against obesity is identical, despite a complete loss of T1INF response. We demonstrated directly that the T1INF response was ablated in these animals, with characteristic response genes completely blunted in their expression in KO mice, and no activation of these genes in response to PSMD9-ASO treatment in WT or KO mice. These findings provide strong evidence that the metabolic improvements observed with PSMD9-ASO treatment is not due to off-target T1IFN activation. PSMD9 functions as a multifunctional protein where suppression of PSMD9 may potentially alter the stability of key metabolic regulators governing adipocyte differentiation, lipid storage or nutrient sensing. Although the precise molecular function of PSMD9 in adipose biology remains incompletely understood, several possibilities warrant further investigation.

Several limitations and caveats exist with the current studies. This includes that all studies were performed only in male mice, and future work should determine whether similar responses occur in females. Second, although our data strongly implicate adipose tissue as the major site of action, additional cell-type-specific genetic models will be required to define the effect of PSMD9-ASO in adipocytes. Third, while indirect calorimetry did not reveal significant alterations in energy expenditure, more comprehensive metabolic flux studies will be necessary to determine whether subtle changes in energy expenditure, mitochondrial function or perhaps futile cycling underlie the reductions in adiposity. Fourth, in the absence of a genetic model of PSMD9 deletion (i.e. KO or floxed model), these ASO studies cannot be recapitulated in PSMD9 null mice, and thus we cannot determine if the ASO effect in lost in those animals. Fifth, longer-term studies will be required to evaluate the potential for the treatment frequency of these ASOs to be extended much longer than once weekly, with current evidence suggesting that ASO treatment in mice could still be effective even at monthly dosing. Lastly, safety and potential of dual therapy with existing obesity therapies, including GLP-1 receptor agonists is an important future study, to understand the potential for these ASOs to act either subsequent to incretin-based therapies, or in conjunction to elicit even greater weight loss effects.

Overall, our findings across multiple preclinical models including different mouse strains, dietary interventions, and treatment paradigms, demonstrate that PSMD9-ASO treatment confers protection against diet-induced obesity. PSMD9-targeting ASOs reproducibly prevented or reduced fat mass accumulation while largely preserving lean mass and improving glucose tolerance when compared with both vehicle and Scr-ASO treated controls. These effects occurred independently of food intake and importantly, the observed reductions in body weight and adiposity were not attributable to the previously reported ASO-mediated activation of the T1IFN response. Instead, our data suggest that these beneficial effects are likely driven by a collection of modest and distinct tissue-specific metabolic adaptations over a sustained period, which collectively promote improved energy balance and metabolic health. For example, this may include SCD1 knockdown in the liver and FAS knockdown in adipose tissue, alongside alterations in other lipid metabolic pathways across tissues, which collectively contribute to the observed reduction in adiposity. Another pathway warranting further investigation is futile cycling. Increased substrate cycling, including triglyceride and fatty acid cycling, or cycles involving lipogenesis and lipolysis, could increase energy expenditure and potentially contribute to the reduced adiposity following PSMD9 suppression. However, these studies remain outside the scope of this first study, and are peripheral to the main aims of this of this study – which have been achieved.

Together, these findings identify this PSMD9-ASO approach as a promising intervention strategy for obesity and its associated comorbidities. Future studies defining the molecular mechanisms downstream of PSMD9-ASO interventions and evaluating combination therapies with existing anti-obesity agents will be important for advancing these findings toward translational studies.

## METHODS

### Animals

All animal experiments were approved by the Alfred Medical Research and Education Precinct (AMREP) Animal Ethics committee and performed in accordance with the ethical guidelines set out by the National Health and Medical Research Council of Australia. Male mice were used for all mouse experiments with access to food (Standard Laboratory Chow SF00-100, High Fat Diet SF04-001, Amylin liver Nash Diet SF18-074: Specialty feeds, Australia) and water *ad libitum*. Mice were group housed with up to 6 mice per cage on a 12hr light/dark cycle at 22 C and ≈50 humidity, weighed weekly, with cages changed weekly. IFNAR KO mice^15^ were a kind gift from Prof Paul Hertzog (Hudson Institute of Medical Research, Melbourne, Australia) and subsequently bred at the AMREP Precinct Animal Centre as heterozygous parental lines to generate a cohort of *Ifnar1* KO mice (IFNAR KO) and littermate IFNAR-WT mice. All mice were bred and sourced (including wild type strains C57BL/6J and DBA/2J) through the AMREP Precinct Animal Centre. All male mice were age and weight matched before being randomly allocated into groups, with respective investigators blinded to saline/ASO treatment and/or genotype. After group allocation, mice were acclimated for 2 weeks prior to the commencement of diet intervention and animal experiments from eight weeks of age. At the completion of each study, mice were fasted for 4–6 h and then euthanised by a lethal dose of ketamine/xylazine and exsanguination, with plasma and tissues collected, weighed, and snap frozen at −80 C.

### Anti-sense Oligonucleotide (ASO) Production and Treatment

All ASOs were designed and synthesised by Ionis Pharmaceuticals, USA, with screening and selection of the lead ASO used, and target sequences of those ASOs having been previously described^7^. Liver targeting PSMD9/Scr-ASOs were identical to native ASOs, except that they were conjugated with the N-Acetylgalactosamine (GalNAc) moiety (Ionis). Whole-body ASOs were diluted in saline to 5mg/ml and GalNAc ASOs (liver targeting) were diluted in saline to 0.2mg/ml. Mice were administered weekly intraperitoneal ASO injections at a dose of 25mg/kg (whole-body) or at 1mg/kg (GalNAc ASOs).

### Whole-Body Energetics, Movement, and Food Intake

Whole-body energetics including energy expenditure (EE), respiratory exchange ratio (RER), food intake, and movement were measured at designated time points using the Promethion High-Definition Behavioural Phenotyping System for Mice (Sable Systems, USA). As previously described, mice were individually housed in metabolic cages for 72hrs, with the first 24hrs used as an acclimation period and the data recorded in the final 48hrs of the experiment used for analysis (12hrs Light/12hrs Dark)^24,25^.

### Body Composition

Body composition in live animals was measured at designated time points using the 4-in-1 EchoMRI (EchoMRI, USA). As previously described, conscious mice were individually placed in the chamber, with lean mass and fat mass recorded as previously described^24,25^.

### Oral Glucose Tolerance Test

Oral glucose tolerance tests (oGTTs) were performed at designated time points at a dose of 2g/kg lean mass, as determined by EchoMRI, after a 5hr fast as previously described^24–26^. Briefly, fasting blood glucose was measured, using a glucometer (Roche, USA) from a sample obtained from the tail vein, before mice received a bolus of glucose (25% in water) via oral gavage. Subsequent blood glucose measures were made at 15, 30, 45, 60, 90, and 120 minutes post-gavage.

### Intraperitoneal Insulin Tolerance Test

Mice underwent an insulin tolerance test after a 4hr fast, at a dose of 0.5U/kg lean mass for chow fed mice and 0.1U/kg lean mass for HFD fed mice, as previously described^25,27^. Initially, fasting blood glucose was measured, using a glucometer (Roche, USA) from a sample obtained from the tail vein, before mice received insulin via an intraperitoneal injection. Subsequent blood glucose measures were made at 15, 30, 45 and 60 minutes post-injection.

### SDS-PAGE and Immunoblot

Tissues were homogenised in radio immunoprecipitation assay buffer (RIPA; Sigma) supplemented with protease and phosphatase inhibitors. Protein quantities were matched and separated by SDS–PAGE, then transferred to PVDF membranes. Membranes were blocked in 3% skim milk for 2 hours and then incubated with primary antibody (see **Supplementary Table 1**) for 16 hours at 4°C. After incubation with primary antibody, membranes were probed with their respective HRP-conjugate secondary anti-mouse or anti-rabbit (Biorad) antibodies in 3% skim milk for 2 hours at room temperature, then visualised with chemiluminescence (Pierce). Approximated molecular weights of proteins were determined from a co-resolved molecular weight standard (BioRad, #1610374). Densitometry analyses were performed in Image Lab (BioRad), with quantification results either normalised to their respective loading control or total protein.

### Plasma Pathology

Fresh animal blood collected via cardiac puncture was centrifuged at 10,000g for 5min at 4 C with 200ul of plasma (supernatant) collected in fresh tubes. Plasma pathology was performed by Vetnostics Pathology (Australia).

### Histology

Liver tissues were fixed in formalin and mounted in paraffin before 4µm sections were cut on a Leica microtome. All sections were stained with Pico Sirius Red (PSR) or haematoxylin and eosin stain (H&E), and slide images were captured using an Olympus Slide scanner VS120 (Olympus, Japan) and viewed in the accompanying software (OlyVIA Build 13771, Olympus, Japan). Fibrotic area quantification was performed as previously described based on threshold analysis in Fiji Image^28,29^. Lipid droplet analysis was performed in QuPath^30^. **Tissue segmentation:** Tissues were identified from the background using a pixel classifier applied to the red channel. The channel was Gaussian-smoothed (sigma of 2) with pixels below a threshold of intensity 220 set as tissue. Annotations of tissue regions were generated with a minimum area of 10,000µm² and a minimum hole area of 1,300µm². Within these tissue annotations, a lipid pixel classifier was applied to the mean RGB intensity. The image was Gaussian-smoothed (sigma of 2) with pixels ≥205 intensity classified as potential lipid regions. Lipid detections were generated with a minimum area of 40µm² and a minimum hole area of 30µm². Shape measurements, namely area and circularity, were added to each detection. **Object classification:** Potential lipid detections were classified by size and then by shape. Objects with circularity ≥0.45 were classified as lipid. The percentage of lipid area was calculated by the sum of the area of the classified lipids detections over the total non-lipid area detected.

### Quantitative PCR (qPCR)

RNA was isolated from tissues using RNAzol reagent and isopropanol precipitation. cDNA was generated from RNA using MMLV reverse transcriptase (Invitrogen) according to the manufacturer’s instructions. qPCR was performed on 10ng of cDNA using the SYBR-green method on a QuantStudio 7 Flex Real-Time PCR System, using primer sets outlined in **Supplementary Table 2**. Primers were designed to span exon-exon junctions and were tested for specificity using BLAST (Basic Local Alignment Search Tool; National Centre for Biotechnology Information). Amplification of a single amplicon was confirmed by melt curve analysis, ensuring an expected temperature dissociation profile was observed. Quantification of a given gene was determined by the relative mRNA level compared with control using the delta-CT method, which was calculated after normalisation to one of two housekeeping genes: *Ppia* or *Rplp0*.

### Transcriptomics

Bulk RNA sequencing was performed as previously described^25^, and raw data is deposited in an online repository. RNA was extracted from gonadal adipose tissue from each group using The Direct-zol RNA MiniPrep kit according to manufacturer’s instructions (Zymo Research). RNA quality for sequencing was determined using the high-sensitivity Agilent RNA ScreenTape Assay for 4200 Tape Station System (Agilent Technologies). Only samples with an RNA Integrity Number (RIN) score >8 were selected for subsequent analysis. DNA libraries were generated as per the manufacturer’s protocol using the NEBNext® Poly(A) mRNA Magnetic Isolation Module kit (New England Biolabs). 1μg total RNA per sample was inputted and adaptor ligation and PCR enrichment was performed using the NEBNext® Multiplex Oligos for Illumina® kit as per manufacturer’s instructions. Libraries for each of the 12 samples were individually bar coded using n6-barcodes and library quantity and quality were assessed as per manufacturer’s instructions using the Agilent D1000 ScreenTape Assay for 4200 TapeStation System. DNA libraries were pooled and then ran on a NovaSeq 6000 sequencer at the Centre of Genomic Medicine at the Alfred Hospital/Monash University.

### Transcriptomics Analysis and Gene Set Enrichment

RNA-sequencing data were processed and analysed similarly to that demonstrated previously ^24,25^. Briefly, quality of raw sequence reads were assessed by FastQC. Optical duplicates were removed using BBMap clumpify (https://github.com/BioInfoTools/BBMap) BBMap version 38.75. The processed reads were aligned to mm10 version of the mouse genome using STAR aligner. Mapped genes were counted using Feature Counts and lowly expressed genes were determined as those that had <1 count per million (CPM) and subsequently removed. PCA was conducted with normalised Log_2_ values to confirm separation of samples based on group, with no significant RNA concentration or read depth effect. Differential expression analysis was performed using DESeq2 and iDEP^31^. Gene Set Enrichment Analysis of upregulated and downregulated gene sets was performed using ShinyGO with Kyoto Encyclopedia of Genes and Genomes (KEGG) and Gene Ontology databases^32^.

### Statistical Analyses

All data were expressed as mean ± standard error of the mean (SEM) unless otherwise stated. Where appropriate, normality was checked using Shapiro-Wilk normality tests. Normally distributed data were analysed by two-way ANOVA (repeated measures in animal studies) with post hoc testing (Fishers LSD) where appropriate, or unpaired Student’s t-test (welch correction). For non-normally distributed data were analysed by Mann-Whitney test. Analyses were performed using PRISM software and a P value of P < 0.05 was considered statistically significant.

### Data Inclusion and Exclusion Criteria

For animal experiments, phenotyping data points were excluded using the following pre-determined criteria: if the animal was unwell at the time of analysis, there were technical issues identified (such as failed beam breaks in Promethion), or data points were identified as outliers using Tukey’s Outlier Detection Method (1.5 IQR < Q1 or 1.5 IQR > Q3). If repeated data points from the same mouse/sample failed QC based on predetermined criteria or several data points were outliers as per Tukey’s rule, the entire animal was excluded from that given analysis (i.e., during glucose tolerance tests, indicating inappropriate gavage). For *in vivo* and *in vitro* tissue and molecular analyses, data points were only excluded if there was a technical failure (i.e., poor RNA quality such as RIN = <7, failed amplification in qPCR, failed injection in mass spectrometer) or the value was biologically improbable. This was performed in a blinded fashion (i.e., on grouped datasets before genotypes were known).

## Supplementary Figure Legends

**Supplementary Figure S1:**
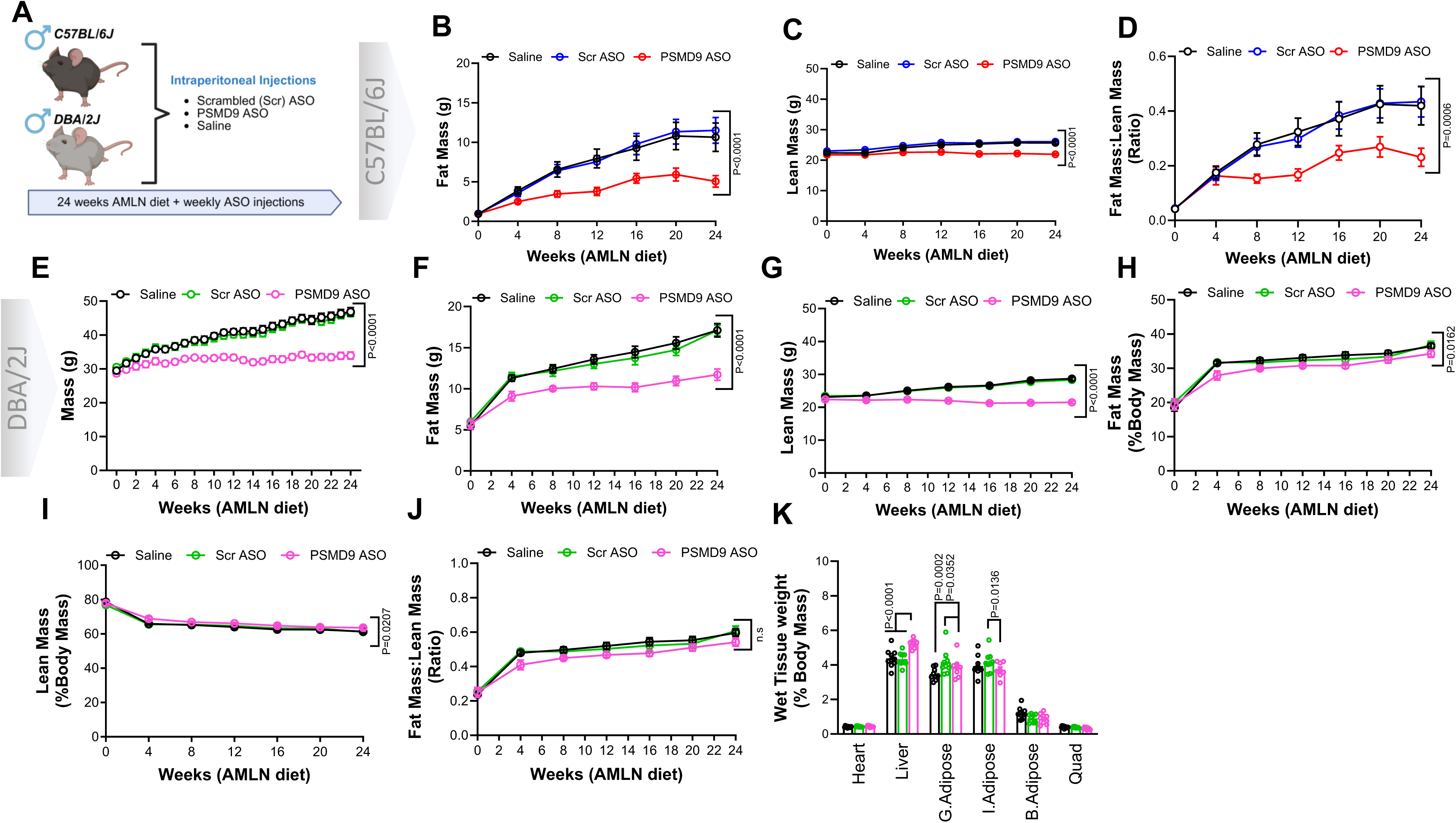
Treatment with ASOs Targeting PSMD9 Protects Against the Development of Obesity. **(A)** Schematic outlining animal study to investigate effect of weekly PSMD9-ASO administration over a 24-week period in C57BL/6J and DBA/2J mice. **(B)** Progressive weekly weight gain in Saline (black, n=12), Scrambled ASO (blue, n=11) and PSMD9-ASO (red, n=12) treated C57BL/6J mice, including body composition; **(C)** lean mass and **(D)** fat to lean mass ratio. **(E)** Progressive weekly weight in DBA/2J mice treated with either Saline (black n=10), Scrambled ASO (green, n=9) or PSMD9-ASO (pink, n=9), including body composition; **(F)** fat mass, **(G)** lean mass, **(H)** percent fat and **(I)** lean mass as a percent of total body mass, **(J)** fat mass to lean mass ratio and **(K)** wet tissue wight as a percent of total body mass. Data are presented as mean ± SEM. P-value determined by two-way ANOVA with correction for multiple comparisons (Panels B-K).

**Supplementary Figure S2:**
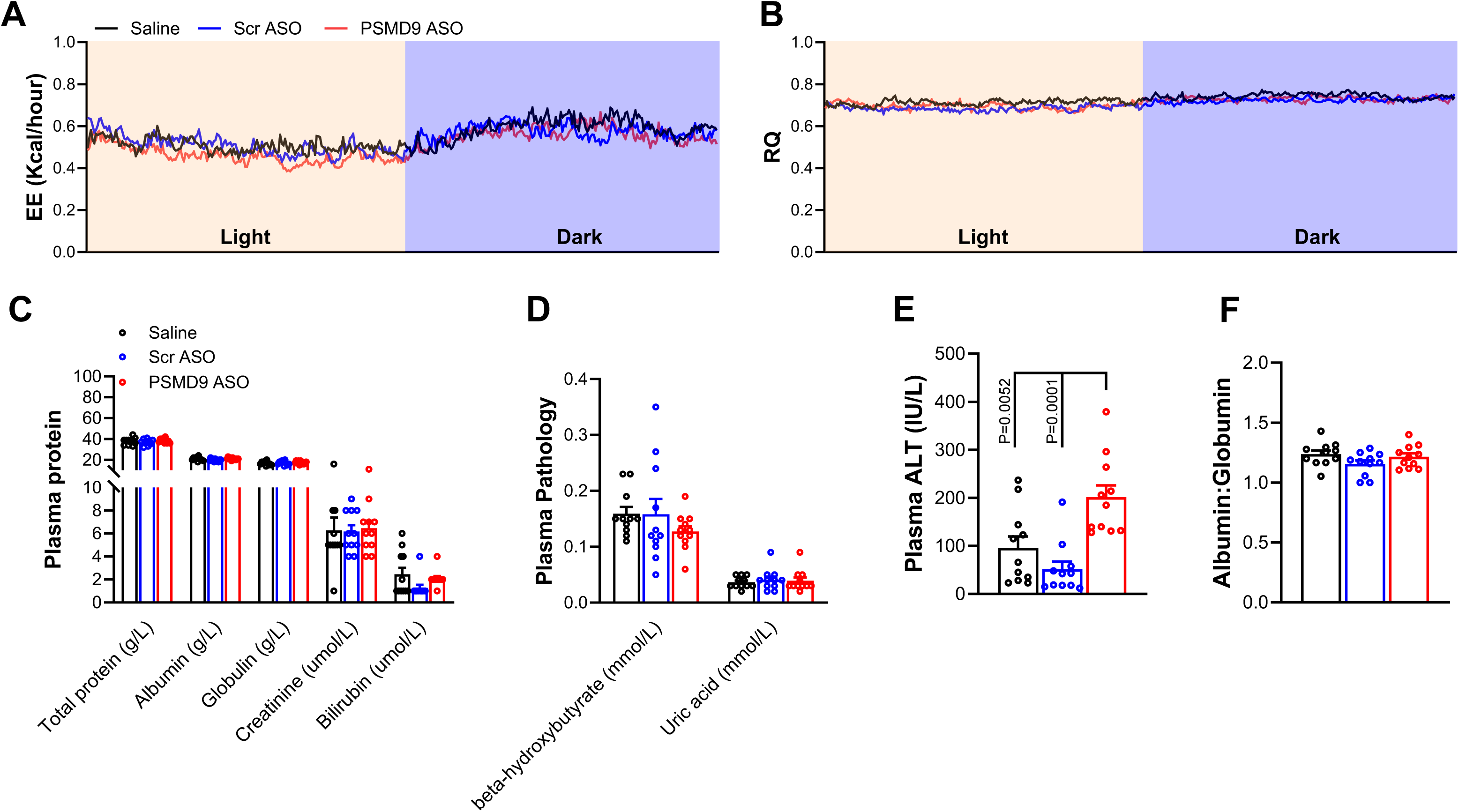
Treatment with whole body PSMD9-ASOs Reduced the Severity of Hepatic Steatosis. C57BL/6J mice treated with either Saline (black, n=12), Scrambled ASO (blue, n=11), or PSMD9-ASO (red, n=12) were placed in metabolic cages (12hrs light & 12hrs dark) to measure **(A)** energy expenditure and **(B)** respiratory quotient. **(C & D)** Plasma pathology was performed to measure and assess systemic metabolic status and toxicity in Saline, Scrambled ASO, and PSMD9-ASO treated C57BL/6J mice, including **(E)** plasma alanine aminotransferase (ALT) and **(F)** albumin to globulin ratio. Data are presented as mean ± SEM. P-value determined by one-way ANOVA with correction for multiple comparisons (panels C-F).

**Supplementary Figure S3:**
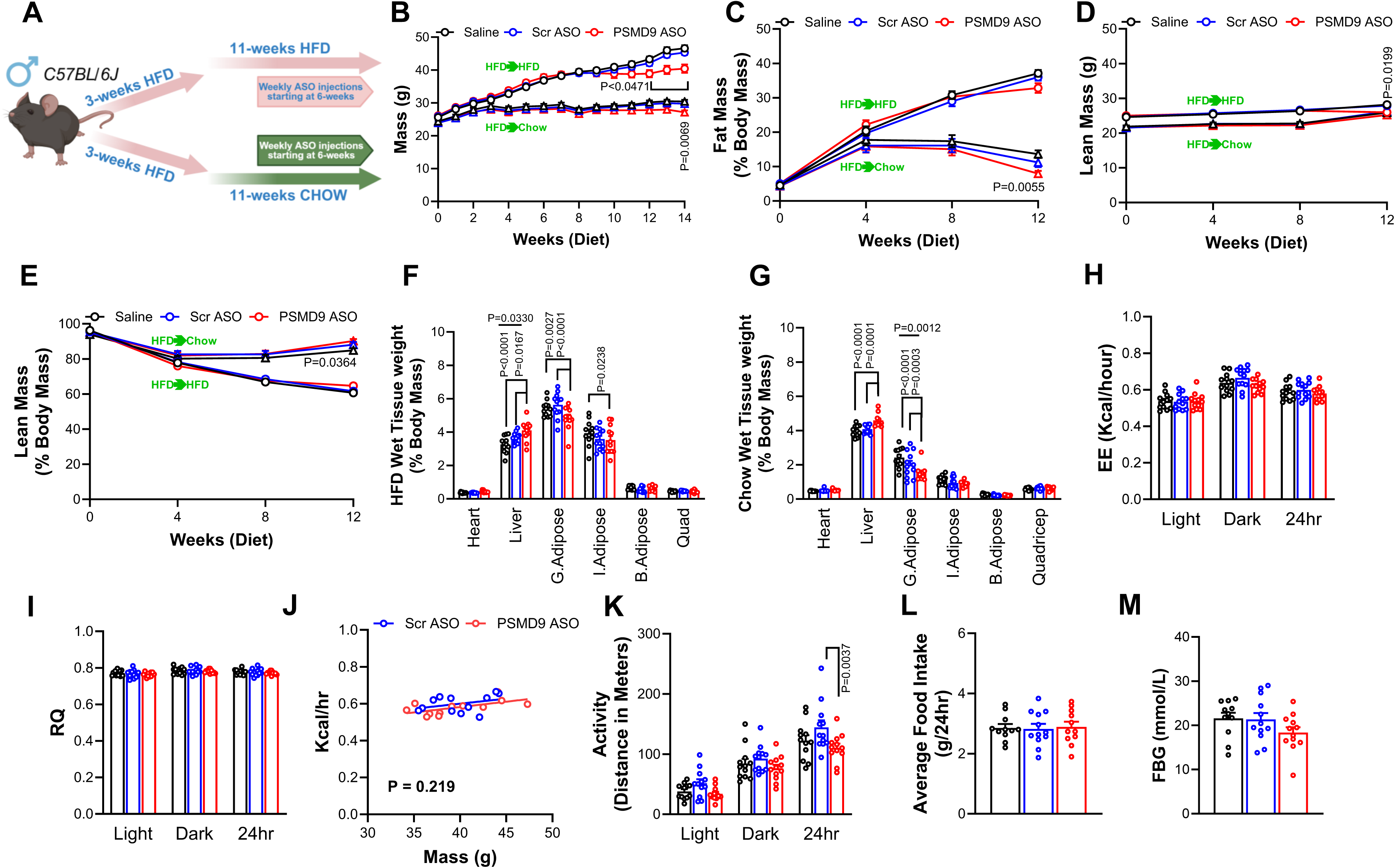
PSMD9-ASOs Improve Adiposity and Glucose Tolerance in Established Obesity. **(A)** Schematic outlining animal study to investigate effect of weekly PSMD9-ASO administration in C57BL/6J mice with established obesity. **(B)** Weekly weight, **(C)** progressive percent fat mass, **(D)** progressive lean mass and **(E)** progressive percent lean mass in C57BL/6J mice with established obesity treated with either high fat diet (HFD) + Saline (black circle, n=12), Scrambled ASO (blue circle, n=12), or PSMD9-ASO (red circle, n=12) or CHOW + Saline (black triangle, n=12), Scrambled ASO (blue triangle, n=12), or PSMD9-ASO (red triangle, n=12). Percent wet tissue weight in ASO treated C57BL/6J mice fed **(F)** HFD and **(G)** CHOW diet. HFD fed C57BL/6J mice treated with either Saline, Scrambled ASO, or PSMD9-ASO were placed in metabolic cages (12hrs light & 12hrs dark) to determine 24hr averages of **(H)** energy expenditure (EE), **(I)** respiratory quotient (RQ), **(J)** EE as a measure of total body mass (ANCOVA), **(K)** total activity and **(L)** food intake. **(M)** Fasting blood glucose measured in HFD + ASO treated C57BL/6J mice after 14-weeks of HFD. Data are presented as mean ± SEM. P-value determined with correction for multiple comparisons by two-way ANOVA (panels B-I & K), or one-way ANOVA (panels L & M) or ANCOVA (panel J).

**Supplementary Figure S4:**
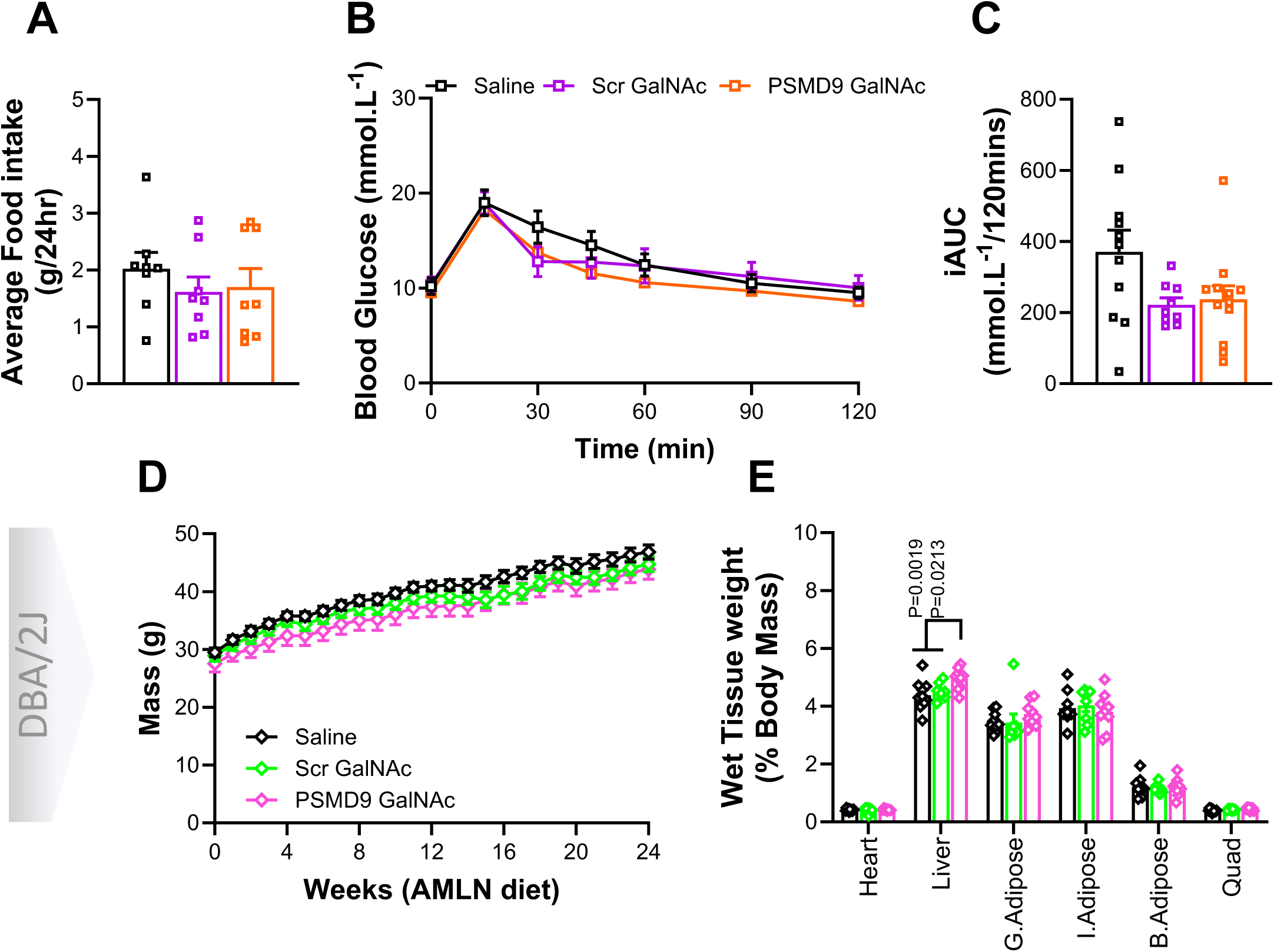
Liver Targeted PSMD9-ASOs do not Impact Body Weight, Adiposity or Metabolic Health. **(A)** average food intake and **(B)** glucose tolerance, with **(C)** corresponding iAUC in C57BL/6J mice treated with either Saline (black, n=12), Scrambled GalNAc ASO (purple, n=12) and PSMD9 GalNAc ASO (orange, n=12). **(D)** Weekly weight and **(E)** wet tissue weight as a percent of total body mass in DBA/2J mice treated with either Saline (black, n=12), Scrambled GalNAc ASO (green, n=12) and PSMD9 GalNAc ASO (pink, n=12). Data are presented as mean ± SEM. P-value determined with correction for multiple comparisons by two-way ANOVA (panels B & D-E) or one-way ANOVA (panels A & C). Saline (black, n=12), Scrambled GalNAc ASO (purple, n=12) and PSMD9 GalNAc ASO (orange, n=12) treated C57BL/6J mice

**Supplementary Figure S5:**
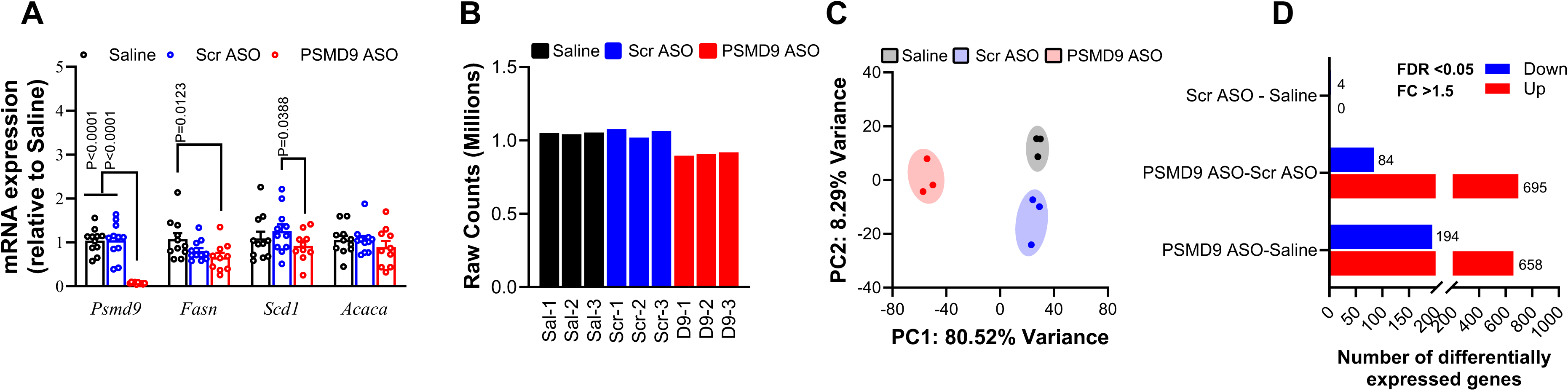
Native PSMD9-ASOs Impart Major Impacts on Adipose Tissue. **(A)** *Psmd9, Fasn, Scd1* and *Acaca* gene expression in gonadal adipose tissue from C57BL/6J mice with established obesity (Figure 3) treated with either high fat diet (HFD) + Saline (black, n=12), Scrambled ASO (blue, n=12), or PSMD9-ASO (red, n=12). Transcriptomic analyses were performed in gonadal adipose tissue from the same C57BL/6J mice with established obesity treated with either high fat diet (HFD) + Saline (black, n=3), Scrambled ASO (blue, n=3), or PSMD9-ASO (red, n=3). Computational analysis of the data was performed including **(B)** comparison of raw counts per mouse, **(C)** principal component analysis (PCA), and **(D)** differential gene expression analyses between all three groups. P-value determined with correction for multiple comparisons by two-way ANOVA (panel A). Analyses of transcriptomic data (B-D) were corrected using Benjamini-Hochberg FDR, n=3/group with DEGs determined by adj P-value < 0.05 and FDR > 1.5.

**Supplementary Figure S6:**
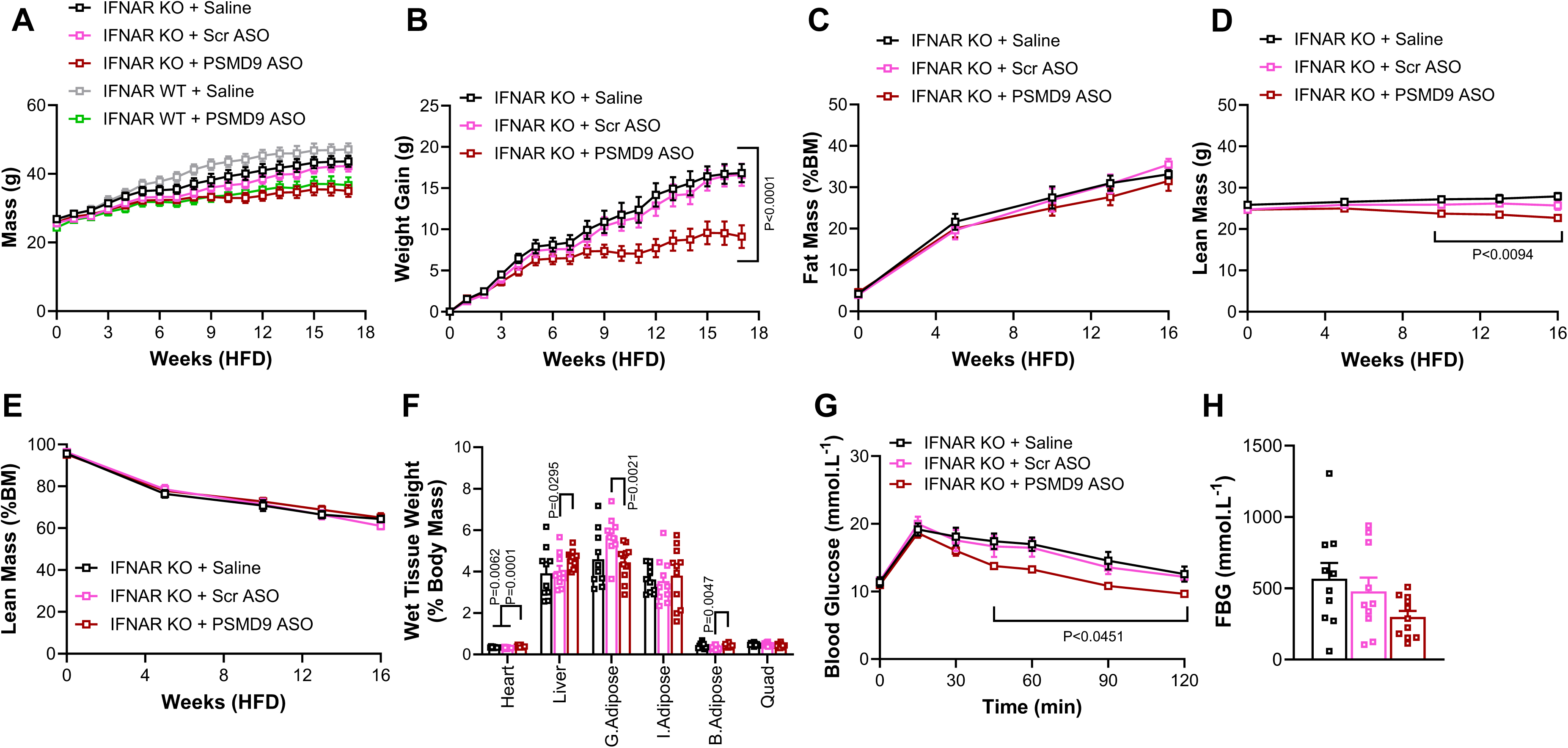
Native PSMD9-ASOs Protect from Obesity in IFNAR-KO Mice. **(A)** Weekly weight in IFNAR WT mice fed a HFD and treated with either Saline (grey square, n=7) or PSMD9-ASO (green square, n=8) and IFNAR KO mice fed a HFD and treated with either Saline (black square, n=9), Scrambled ASO (pink square, n=10), or PSMD9-ASO (red square, n=11). **(B)** weight gain, **(C)** monthly percent fat mass, **(D)** monthly lean mass, **(E)** monthly percent lean mass and **(F)** wet tissue weight as a percent of total body mass in IFNAR KO mice fed a HFD and treated with either Saline (black square, n=9), Scrambled ASO (pink square, n=10), or PSMD9-ASO (red square, n=11). **(G)** Glucose tolerance and **(H)** fasting blood glucose in ASO treated IFNAR KO mice. Data are presented as mean ± SEM. P-value determined with correction for multiple comparisons by two-way ANOVA (panels A-G) or one-way ANOVA (panel H).

## DATA AVAILABILITY

All raw and processed data is, and will be made available, in supplemental data and tables and in online repositories where appropriate.

## ACKNOWLEDGMENTS

We acknowledge the administrative and technical support from all members of the MMA Laboratory at the Baker Heart & Diabetes Institute, particularly past alumni including Medini Reddy, Morgan Griffiths, Eleanor Gould and Tim Sikora. We thank Dr Richard Lee, previously from Ionis Therapeutics, for prior support with ASO design and synthesis. We are grateful to Prof Paul Hertzog from the Hudson Institute of Medical Research, Melbourne, Australia for providing us with the IFNAR1 KO mice. We also thank Thomas de Aguiar Vallim (UCLA and Elizabeth Tarling (UCLA), ongoing collaborations relating to this work. We thank the Monash Histology Platform, Monash University, Melbourne, Australia for help with liver histology, and the Metabolomics Laboratory/Platform at Baker Heart & Diabetes Institute for expertise in lipid profiling.

## AUTHOR CONTRIBUTION

BGD and ACC conceived the study and co-designed the experiments with STB and MFK. BGD, STB and MFK wrote and edited the original manuscript. STB, MFK, CY, YL, and KHL performed animal experiments and phenotyping. STB, MFK, BGD, ACC, CY, YL, KHL, ACP, APN, SRC, and MI either analysed data, processed tissue samples, or performed molecular, biochemical and bioinformatics analyses. BGD and ACC provided reagents, experimental advice, and access to infrastructure and resources. All authors read and approved the manuscript.

